# The language network responds stronger to connected than to unconnected sentences

**DOI:** 10.64898/2026.11.20.768057

**Authors:** Fiona Paliga, Zenab Javed, Manuel Unnerstall, Shaza Haj Mohamad, Andreas Neef, Julia My Van Kube, Vesna Malinova, Tammam Abboud, Peter Dechent, Christian H. Riedel, Nicole E. Neef

## Abstract

**Purpose:** Reliable functional mapping of the language network and preoperative assessment of language lateralization are essential for safe and effective craniotomy in patients with lesions in language-eloquent brain regions, yet existing fMRI language paradigms are often time-consuming and place substantial cognitive and economic demands on patients and diagnostic services.

**Methods:** We therefore tested ecologically valid German stimulus sets designed to optimize measurement efficiency. Forty-three healthy participants (mean age = 38 years, range = 20 - 64; 24 women) were scanned on a 3T MRI while reading or listening to 4-, 8-, and 12-word sentences and pseudoword lists or texts, each followed by a decision task. Sentences were selected from German literary classics and text passages from general-interest science magazines and matched across conditions for lexical and structural properties.

**Results:** Region-of-interest analyses in subject-specific, functionally defined language-sensitive regions showed robust activation of the left-hemispheric language network across modalities. Within participants, texts elicited significantly larger effect sizes than sentences, and longer sentences generally produced stronger responses than shorter ones. Task manipulations expected to increase cognitive demand (short 4-word sentences with frequent probes, coherent texts requiring deeper semantic integration) did not enhance responses in the domain-general multiple demand network, indicating that executive-control activity did not overshadow language-selective activation, whereas substantial overlap between default mode and language-sensitive regions in the left temporal lobe was consistent with relatively automatic processing of connected language.

**Conclusions:** These findings indicate that ecologically valid sentence- and text-based paradigms can provide efficient, language-specific fMRI protocols for preoperative diagnostics and individualized presurgical planning in patients with lesions near language-eloquent cortex.

## INTRODUCTION

For patients with language-eloquent pathologies, presurgical planning relies on noninvasive language mapping to support the onco-functional balance between maximal resection and preservation of language function [1,2]. This need has driven the widespread adoption of noninvasive language mapping and tractography [3–6] in addition to intraoperative direct electrical stimulation [7,8]. However, conventional task-based fMRI paradigms often show limited specificity, in the sense that activation in cortical language regions does not reliably predict long-term postoperative language outcomes, pointing to a “causality gap” between task-related activation and functionally essential language sites [9]. Extensive work in healthy adults has characterized the functional organization of the language network and its relationship to domain-general systems, largely with the goal of dissecting the underlying cognitive architecture rather than optimizing clinical protocols [10]. By contrast, preoperative language fMRI requires paradigms that yield robust, reliable, and time-efficient activation of language-selective cortex at the level of individual patients, suitable for use as practical localizers in the clinical setting [6,11,12]. Bridging this gap between basic and clinical work calls for systematic evaluations of candidate language tasks in healthy participants, explicitly targeting signal strength, selectivity, and reliability under constraints that are realistic for presurgical mapping.

A critical but underexplored determinant of language-network activation is the degree of linguistic connectedness of the stimulus material, that is, the extent to which successive words and sentences form a coherent, information-rich sequence of content rather than isolated fragments. While sentences are known to robustly engage language-selective regions [13,14], many clinical paradigms rely on relatively short or weakly connected linguistic input [6,11,12,15]. These may drive the core language system only suboptimally, and thus limit signal strength and render activation patterns more susceptible to domain-general co-activation. In particular, task-based paradigms can recruit the multiple demand (MD) and default mode (DM) networks as a function of task difficulty, structure, and attentional demands [16,17], making it unclear whether more connected language input primarily strengthens responses within language-selective cortex or instead draws more heavily on MD and DM resources. To improve the interpretability and clinical utility of preoperative language fMRI it is therefore essential to optimize stimulus design in a way that enhances language-selective responses by increasing informational connectedness and coherence while minimizing domain-general engagement.

Here, we systematically test the hypothesis that the language network responds more strongly to connected than to less connected linguistic input. To this end, we selected German stimuli from natural sources, including literary classics and National Geographic text passages, that varied in informational connectedness and length, comprising 4-, 8-, and 12-word sentences, short coherent texts, and matched pseudoword lists. Converging evidence suggests that core language regions show highly overlapping responses to spoken and written sentences, consistent with a largely supramodal organization of sentence processing [18–20]. At the same time, studies also report modality-specific differences in activation strength and in the recruitment of adjacent sensory and association areas, indicating that language-selective cortex is embedded within broader, modality-sensitive networks [21]. In addition to reading, we therefore also tested listening, both to exploit the supramodal component of the language network and to capture any modality-dependent differences that may be relevant for clinical task selection in patients who can only be tested auditorily. Forty-three healthy participants underwent fMRI during reading and listening, and activation was quantified using group-constrained subject-specific fROIs [14] in the language, MD, and DM networks. By comparing connected and less connected stimuli across modalities, we aim to identify stimulus conditions that yield strong, selective, and clinically efficient language-network activation.

## METHODS

### Participants

We tested 43 healthy volunteers (24 females) between the ages of 20 and 64 years (mean age: 38 years). Exclusion criteria were contraindications for MRI, as well as underlying neurological or psychiatric conditions, chronic illnesses requiring medication, history of substance abuse, and current pregnancy or breastfeeding. All subjects were native speakers of German and had normal or corrected-to-normal vision. Based on the Edinburgh Handedness Inventory [22], 40 participants were right-handed, one was ambidextrous and two were left-handed. The study was approved by the Ethics committee of the University [blinded for review] and conducted in accordance with the guidelines of the Declaration of Helsinki. All participants gave written informed consent and received financial compensation for their participation.

### Language Mapping Paradigm

German 4-, 8-, and 12-word sentences and short texts were selected from the Gutenberg library and National Geographic text passages, and matched lists of pronounceable nonwords were constructed for each trial. Blocks followed a blocked design (Fig. 1a) and consisted of either five 4-word sentences, three 8- or 12-word sentences, or one text trial, paired with corresponding nonword lists. To approximately equalize the total number of words across conditions, the number of trials per run depended on the condition: a single run comprised 45 4-word sentences, 21 8-word sentences, 15 12-word sentences, or 4 text trials and 21 8-word sentences, together with the respective matched nonword lists. This yielded a total of 180 words per run in the 4- and 12-word conditions, 168 words in the 8-word condition, and 184 words in the text condition, so that the overall number of presented words was broadly comparable across conditions. Nonwords were matched to their sentence or text counterparts for number of syllables, phonetic complexity, and phonotactic structure and were generated with the software Wuggy [23], a multilingual pseudoword generator that matches pseudowords to word templates in subsyllabic structure and phonotactic constraints.

For reading, stimuli were presented visually on a back-projection screen located at the rear of the scanner bore and illuminated by a video projector positioned outside the magnet. Participants viewed the stimuli via a mirror attached to the head coil and were instructed to read the words silently. Each word of a sentence, text, or nonword list was presented sequentially, each trial was followed by a forced-choice recognition probe in which participants indicated which of two displayed words or nonwords had appeared in the current trial (example trials are shown in Fig. 1b). Responses were collected via an MRI-compatible button box, with participants using their left and right hands to select the respective response options.

For listening runs, all sentence, text, and nonword stimuli were converted to audio files using a text-to-speech engine (NaturalReader, NaturalSoft Ltd., https://www.naturalreaders.com/) with a neutral female voice and standardized speech rate. The resulting sound files were delivered via MRI-compatible earbuds (MagnaCoil Tim4G headphones, Siemens Healthineers) equipped with disposable earplugs (MagnaPlugs) providing approximately 30 dB noise suppression, and volume was adjusted individually to ensure comfortable audibility. Outside the scanner room, a Windows computer running the software Presentation (Version 23.0, Neurobehavioral Systems, https://www.neurobs.com) controlled stimulus delivery to the projector and headphones.

The four different experimental runs differed in the combination of sentence-length and text conditions included in each session, as summarized in Fig. 1c. Each variant of runs lasted between 11 and 18 minutes. During each session, one or two runs from a chosen variant were acquired (see Supplementary Table 1), and the order of conditions and blocks was pseudorandomized for each participant to account for order effects. Before entering the scanner, participants practiced the task with a separate set of stimuli to familiarize themselves with the procedure.

**Fig. 1.**
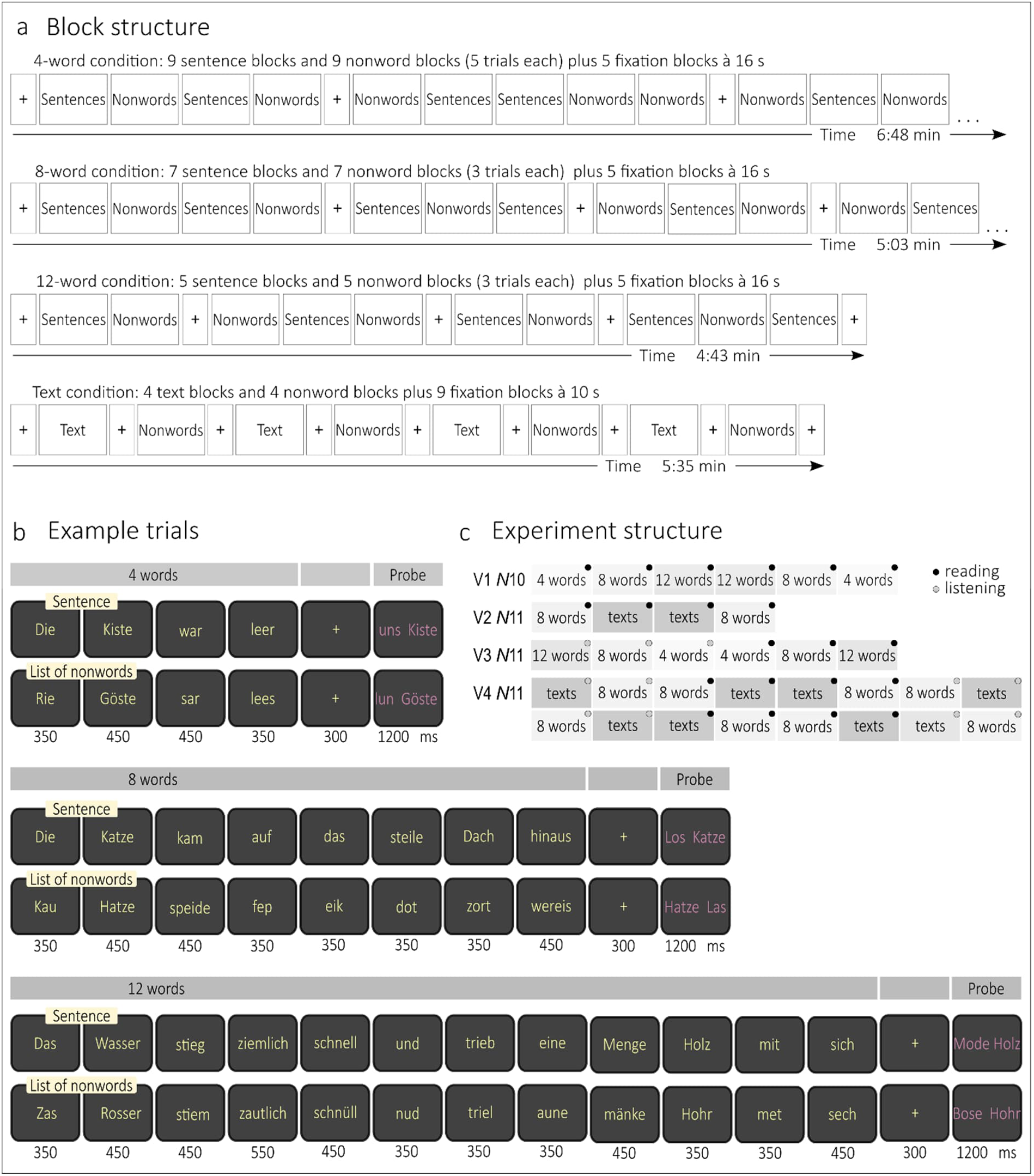
Language mapping tasks. (a) The experimental conditions were presented in blocks of language and nonword stimuli, interspersed with resting periods. Depending on sentence length, blocks contained either 5 (short) or 3 (long) sentence trials, whereas each text was presented in a single block. were approximately matched across conditions, fewer trials were shown in the long-sentence condition and overall presentation time was shorter. (b) Example trials from the sentence conditions with matched nonword lists. The duration of word presentation depended on word length. Words with up to 4 letters were shown for 350 ms, words with up to 7 letters for 450 ms, and longer words for 550 ms. (c) Structure and number (N) of participants for each variant (V) of experimental runs.

### Data Acquisition

MRI data were acquired using a 64-channel, phased-array head coil in a 3 Tesla MAGNETOM Prisma Fit at University Medical Center Göttingen. The participants were examined in a supine position. Sagittal T1-weighted structural data were acquired with a 3D turbo fast low-angle shot sequence with TR = 1900 ms, TE = 2.26 ms, TI = 900 ms, flip angle = 9°, 256 mm FoV, and 7/8 Fourier phase encoding as whole-brain anatomical reference data at a spatial resolution of 1 × 1 × 1 mm³, voxel size. For functional MRI, a gradient-echo echo-planar imaging sequence (TR = 1500 ms, TE = 30 ms, flip angle = 72°, parallel acquisition factor 2, simultaneous multi-slice factor = 3, FoV = 208 mm, 63 slices) was used with an in-plane resolution of 2 × 2 mm², a distance factor of 10% and a 104 × 104 acquisition matrix. We acquired between 373 and 685 volumes per run.

### MRI data analyses

MRI data was processed using FEAT within the FMRIB Software Library v6.0 [24–26]. For preprocessing, the fMRI data was registered to the T1-weighted structural image as anatomical reference, which we previously segmented using FastSurfer [27]. Then, the data was registered into standard space using the template MNI152 T1-weighted structural image. We further performed motion correction, spatial smoothing with a 5 mm full-width half-maximum Gaussian kernel to reduce noise, intensity normalization, and temporal filtering using a high pass filter with a cut-off time of 90 s to reduce artefacts. For the first-level analyses General Linear Modelling was applied to generate the individual contrasts for sentences/texts vs. nonword lists. Convolution of word and pseudoword stimulation periods with the hemodynamic response function was performed applying the gamma variate and including the temporal derivative. The resulting sentences > nonwords and text > nonwords contrast maps were thresholded at *Z* = 3.1 with a cluster probability threshold of *p* = 0,05.

Next, a group-constrained subject-specific analysis was performed using parcels to select individual fROIs for the sentences > nonwords and texts > nonwords contrast in the language, MD, and DM network. The language network mainly includes perisylvian cortex in temporal, frontal and parietal lobes, forming a network which is left-lateralized in most humans [28–32], parcels from [14]. The MD network recruits regions from the medial and superior fronto-parietal axis together with the cingulate gyrus and anterior insula [33,34], parcels from [35]. The DM network recruits medial the prefrontal cortex, cingulate gyrus and precuneus, together with lateral parietal and temporal regions and the hippocampus [36–38], parcels from [39,40].

As there are interindividual neuroanatomical differences in language processing, this subject-specific approach provides a more precise and functionally tailored characterization of language-related brain activity [14,41]. The fROIs were defined for each of the two runs per subject per condition. If only one run for each of the three sentence conditions was measured, three higher-level FEAT analyses were initially carried out aggregating data from two of the three conditions. In this case, fROIs were determined using the combined data of two sentence conditions. To define subject-specific fROIs, voxels within each parcel were ranked according to their t-values for the sentences > nonwords and text > nonwords contrast, and the top 10% were selected as the fROI. The fROIs were then applied as masks to the fMRI data from the other run of the same condition, or, when combined data of two sentence conditions were used for fROI determination, to the run of the left-out condition. This masked data from the other run was used to extract effect sizes for the haemodynamic response within each fROI. This method ensured that fROI definition and response estimation were based on independent data preventing circularity [42].

Additionally, the overlap of the fROIs was calculated within subjects between both runs of the same condition and between runs of the same condition in different subjects. For this, activation maps including overlapping voxels between two runs of the same condition within-subject and between-subjects were generated. Then, the overlap was quantified using the dice coefficient, defined as (2(A∩B))/(A+B) [43] with A and B representing the number of voxels in the fROIs of the two runs.

### Statistical analyses

Statistical analyses were performed in RStudio version 2025.09.02 with R version 4.4.3, using linear mixed-effects models (*nlme* package) and estimated marginal means were obtained with the *emmeans* package.

Our primary question was whether the type of language material used to stimulate the language network (short sentences, long sentences, or continuous texts) influences the strength of the neural response. To address this, we modelled effect sizes as a function of language material while controlling for imaging-related factors (modality and region). We first averaged repeated runs within each subject and condition, so that the dependent variable was a single effect size per subject per condition. To control for imaging-related variation, modality and region and their interaction (modality × region) were included as fixed effects in a baseline model. The primary predictor of interest, language material (4-word, 8-word, 12-word sentences, texts), was then added as a fixed effect to the baseline model. To test whether the effect of material varied across regions, we successively included material × region, material × modality, and material × region × modality interaction terms. Random intercepts for subject and experiment accounted for within-subject dependence and systematic differences across experiments. Model comparisons were performed using likelihood-ratio tests under maximum-likelihood estimation, comparing the baseline model (without material) to models that incrementally added main and interaction effects of material (material; material × region; material × modality; material × region × modality). When a significant interaction was detected, we obtained estimated marginal means for each level of material within the relevant factors and conducted Bonferroni-corrected pairwise comparisons.

**Fig. 2.**
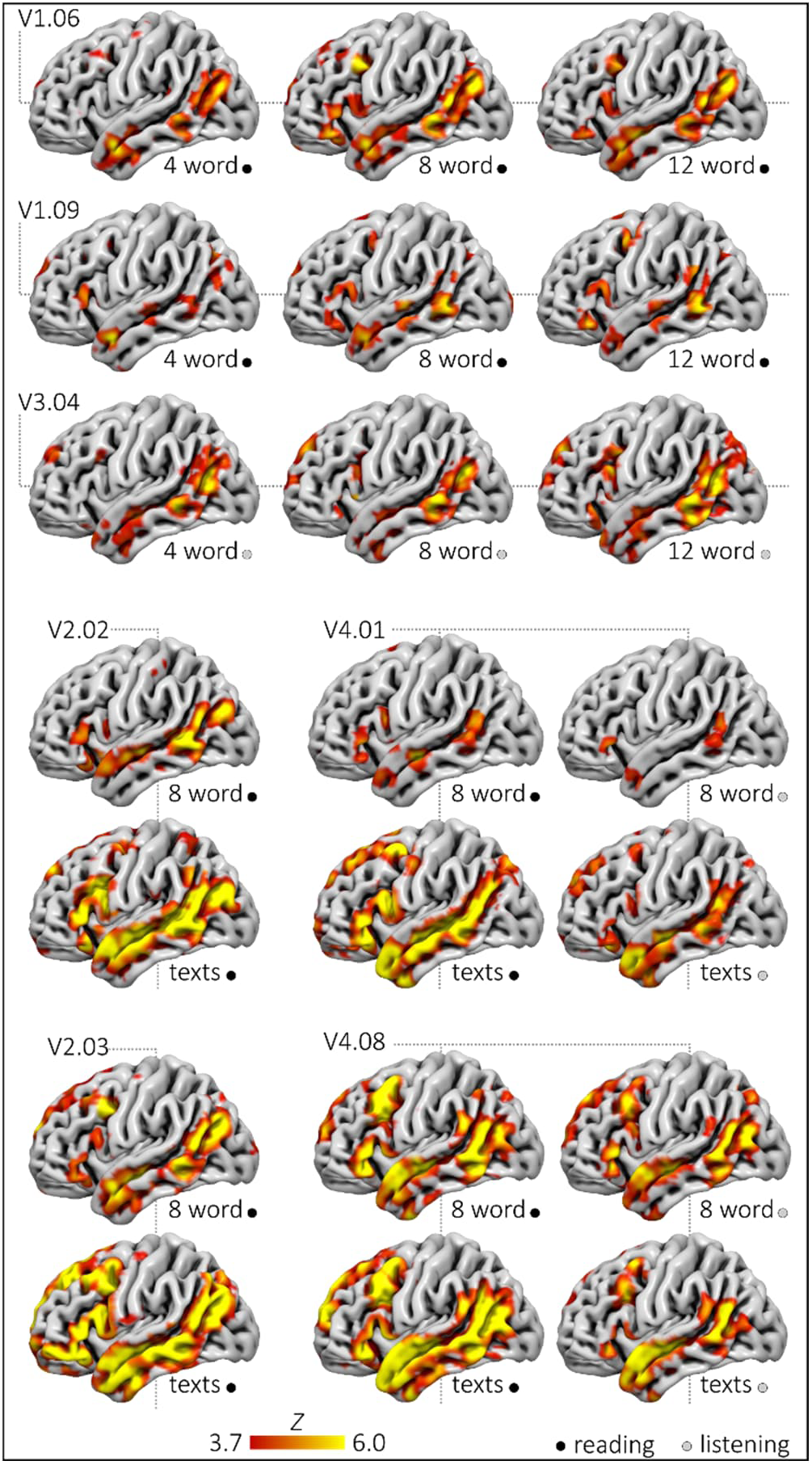
Individual left-hemispheric activation maps for sentences > nonwords and texts > nonwords during reading and listening in seven example participants. Statistical maps were obtained using a higher-level fixed-effects analysis (FSL FEAT). Contrast maps are cluster-corrected with a cluster-forming threshold of Z > 3.1 and a cluster-level significance of p<.05; for visualization, maps are displayed at Z > 3.7.

We also examined, whether the conditions elicited similar or different voxels in the language-related cortical partitions. To this end, we computed dice coefficients (DCs) to quantify the spatial overlap of the activations. Within-subject DCs were calculated by comparing two runs of the same language material and modality within each participant, whereas between-subject DCs were calculated by comparing runs of the same language material and modality across the different participants. For the between-subject analysis, DCs were averaged across both runs within each condition, yielding a single DC per condition. We then modelled DCs as a function of comparison type (within-subject vs. between-subject) while controlling for task-related factors (modality and material). Modality, material and their interaction (modality × material) were included as fixed effects in a baseline model, to which comparison type was added as the primary predictor of interest. To assess whether the effect of comparison type was influenced by language material or modality, we successively added comparison type × material, comparison type × modality, and comparison type × material × modality interaction terms. Random intercepts for subject and experiment accounted for within-subject dependence and systematic differences across experiments. Models were compared using likelihood-ratio tests under maximum-likelihood estimation, contrasting the baseline model (without comparison type) with models that incrementally added the main and interaction effects of comparison type. When a significant interaction was detected, we obtained estimated marginal means for each level of comparison within the relevant factor(s) and conducted Bonferroni-corrected pairwise t-tests.

A subsequent question was whether texts and sentences activate similar regions within the language network. For each subject and modality, we first generated maps containing only top10% voxels that were active in both runs of the 8-word-sentence condition and, separately, maps containing only top10% voxels that were active in both runs of the text condition. Using these two overlap maps, we then calculated DCs per subject and modality, quantifying the spatial overlap between sentence- and text-evoked activation.

Finally, we tested whether neural responses to the language localizer were greater in the language network than in the MD and DM networks, as an index of preferential engagement of the language network. To reduce model complexity, effect sizes were first averaged across runs and regions within each subject, condition, and network, and then normalized to those of the language network within each subject and condition. This yielded a single relative value per subject and condition, reflecting the ratio effect size for the MD and DM networks, with the language network fixed at 1 by definition. Effect sizes were modelled as a function of brain network, controlling for material and modality. The baseline model included modality, material, and their interaction. Brain network (language, MD, DM) was then added as the main predictor, along with its interactions with modality and material, including the three-way interaction. Random intercepts for subject and experiment accounted for within-subject dependence and between-experiment variability. Models were compared using likelihood-ratio tests under maximum likelihood estimation. When significant interactions emerged, estimated marginal means were computed and Bonferroni-corrected pairwise comparisons were conducted.

## RESULTS

For the sentence- and text-level contrasts (sentences > nonwords and texts > nonwords), individual activation maps indicated that, across modalities, experimental conditions engaged the core cortical regions of the left-hemispheric language network, including the superior temporal sulcus and middle temporal gyrus across both anterior and posterior temporal regions, as well as the inferior frontal gyrus and posterior middle temporal gyrus (Fig. 2). Substantial inter-individual variability was observed. In some participants, short sentence conditions were associated with less consistent activation in frontal regions.

### Language material modulates activation in the language network

Adding language material as a fixed effect significantly improved model fit compared with the baseline model including only modality, region, and their interaction (Model 1 vs. Model 2: *χ*²(3) = 209.4, *p* < 0.001). This indicates that the type of language material significantly influenced activation strength in the language network. Extending the model with a material × region interaction provided an additional improvement in fit (Model 2 vs. Model 3: *χ*²(15) = 65.7, *p* < 0.001), indicating that the effect of language material varied across regions. Including the material × modality interaction also yielded a significant improvement in fit (Model 3 vs. Model 4: *χ*²(15) = 20.4, *p* < 0.001), suggesting that material effects differed between modalities. By contrast, adding the three-way material × region × modality interaction did not improve model fit (Model 4 vs. Model 5: *χ*²(15) = 9.7, *p* = 0.836), indicating no evidence for a higher-order interaction between material, region, and modality.

Pairwise comparisons between language material levels within each region were obtained from the linear mixed-effects model using estimated marginal means, with Bonferroni correction for six comparisons per region and denominator degrees of freedom based on the mixed-model “containment” method. Post-hoc comparisons of language material within each left-hemispheric language region, averaged across reading and listening, showed that texts elicited the largest effect sizes, followed by 12- and 8-word sentences, with 4-word sentences producing the smallest responses. For example, in LeftPostTemp, texts produced significantly larger effect sizes than 12-word (*t*(882) = 4.1, *p* < 0.001), 8-word (*t*(882) = 6.1, *p* < 0.001), and 4 word sentences (*t*(882) = 7.5, *p* < 0.001) and 8- and 12-word sentences both exceeded 4-word sentences (12-word-4-word: *t*(882) = 3.5, *p* = 0.003, 8-word-4-word: *t*(882) = 2.9, *p* = 0.022), while 12- and 8-word sentences did not differ significantly from each other (*t*(882) = 1.1, *p* = 1.0, Bonferroni-adjusted). A similar pattern was observed in LeftAntTemp and LeftIFGorb; in LeftIFG the difference between 12- and 4-word sentences was not significant, in LeftMFG 8- and 4-word sentences did not differ, and in LeftMFG no differences were found between sentence conditions, whereas the text condition consistently produced larger responses than all sentence conditions in every region. Fig. 3 illustrates the condition-specific effect sizes for the six language-related regions of the left hemisphere, and Supplementary Table 2 reports the detailed results of the pairwise post-hoc comparisons.

When these pairwise differences were examined separately for each modality, the reading condition largely reproduced the pattern described above, especially the stronger responses to texts compared with sentences (Supplementary Table 3). During listening, this pattern was attenuated and restricted to the temporal and parietal language-related regions (Supplementary Table 4).

**Fig. 3.**
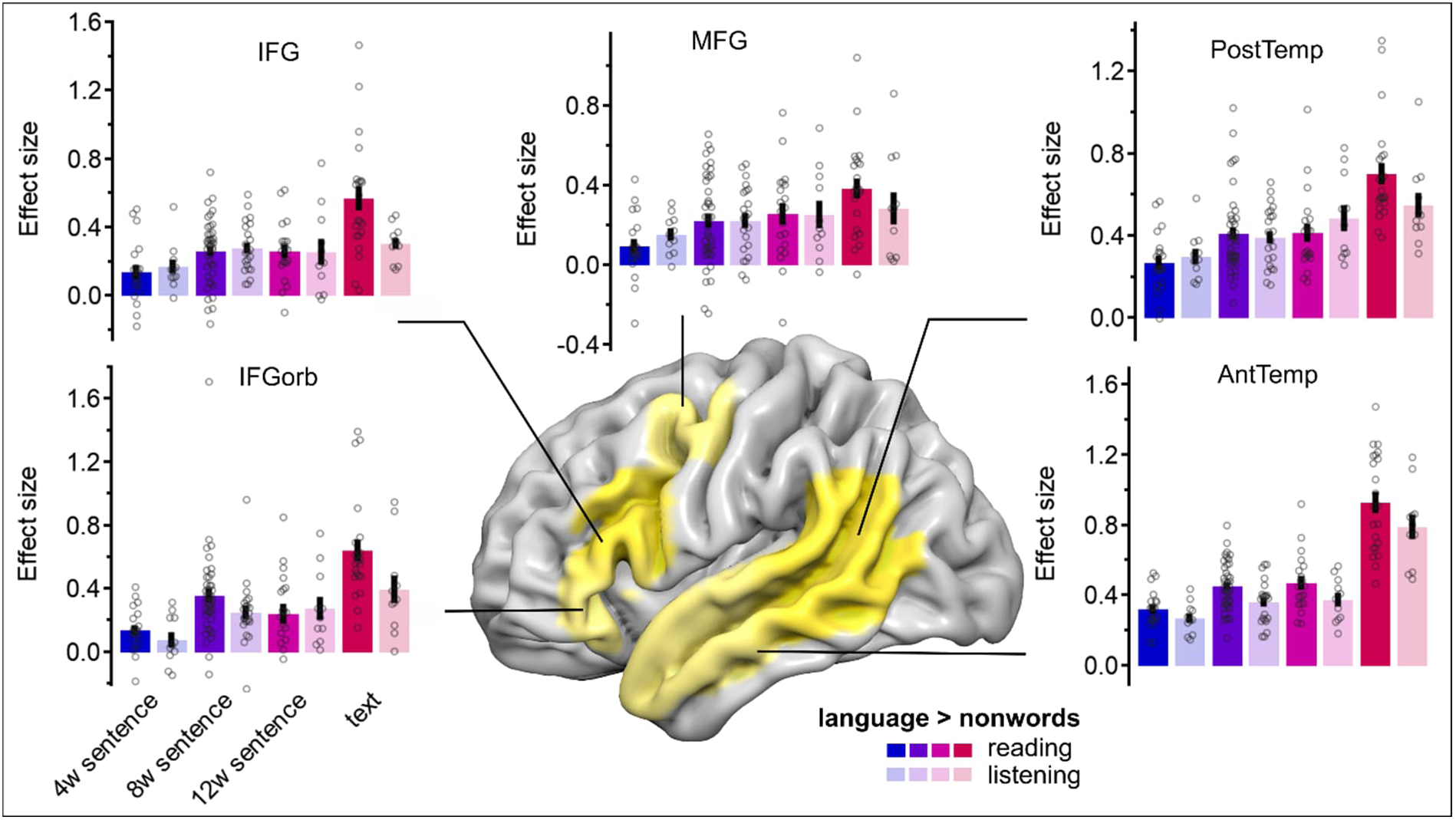
The language network shows stronger responses to connected than to unconnected sentences. Effect sizes are shown for each condition within the language parcels in the left hemisphere. Error bars represent the standard error of the mean.

### Subject-specific activation patterns in the language network

We observed interindividual differences in the localization of language activation. This was examined by modelling DCs as a function of comparison type (within-subject vs. between-subjects). Adding comparison type as a fixed effect significantly improved model fit compared with the baseline model including only modality, material, and their interaction (Model 1 vs. Model 2: *χ*²(1) = 487.2, *p* < 0.001). This indicates that the type of comparison significantly influenced the spatial similarity of the activation pattern of the top 10% voxels. Extending the model with a comparison type × material interaction provided an additional improvement in fit (Model 2 vs. Model 3: *χ*²(3) = 33.9, *p* < 0.001), indicating that the influence of comparison type varied across language materials. However, adding the comparison type × modality interaction did not improve model fit (Model 3 vs. Model 4: *χ*²(1) = 0.4, *p* = 0.509), indicating no evidence for an interaction of comparison type and modality. Pairwise t-tests between within-subject- and between-subjects-comparison were performed for each language material using estimated marginal means from the linear mixed-effects model, with denominator degrees of freedom based on the “containment” method. The results showed that DCs within-subjects were significantly larger than DCs between-subjects for all language materials including texts with *t*(201) = 26.9, *p* < 0.001 (*M*_within_ = 0.613, 95% CI [0.584;0.642], *M*_between_ = 0.154, 95% CI [0.138;0.170]), 12-word-sentences with *t*(201) = 12.3, *p* < 0.001 (*M*_within_ = 0.458, 95% CI [0.400;0.517], *M*_between_ = 0.139, 95% CI [0.123;0.154]), 8-word-sentences with *t*(201) = 27.9, p < 0.001 (*M*_within_ = 0.541, 95% CI [0.510;0.572], *M*_between_ = 0.152, 95% CI [0.140;0.164]), and 4-word-sentences with *t*(201) = 11.6, *p* < 0.001 (*M*_within_ = 0.440, 95% CI [0.376;0.503], *M*_between_ = 0.136, 95% CI [0.117;0.154]), see Fig. 4a and Supplementary Table 5.

Sentences and texts activated similar language-sensitive regions. Within-subject DCs comparing sentence and text conditions were generally large, with a DC of 0.54 (95% CI [0.51; 0.58]) for reading and 0.48 (95% CI [0.43; 0.53]) for listening, indicating substantial spatial overlap between sentence- and text-evoked activation (Fig. 4b), with no significant difference between listening and reading (*t*(21.09) = -1.96, *p* = 0.063).

**Fig. 4.**
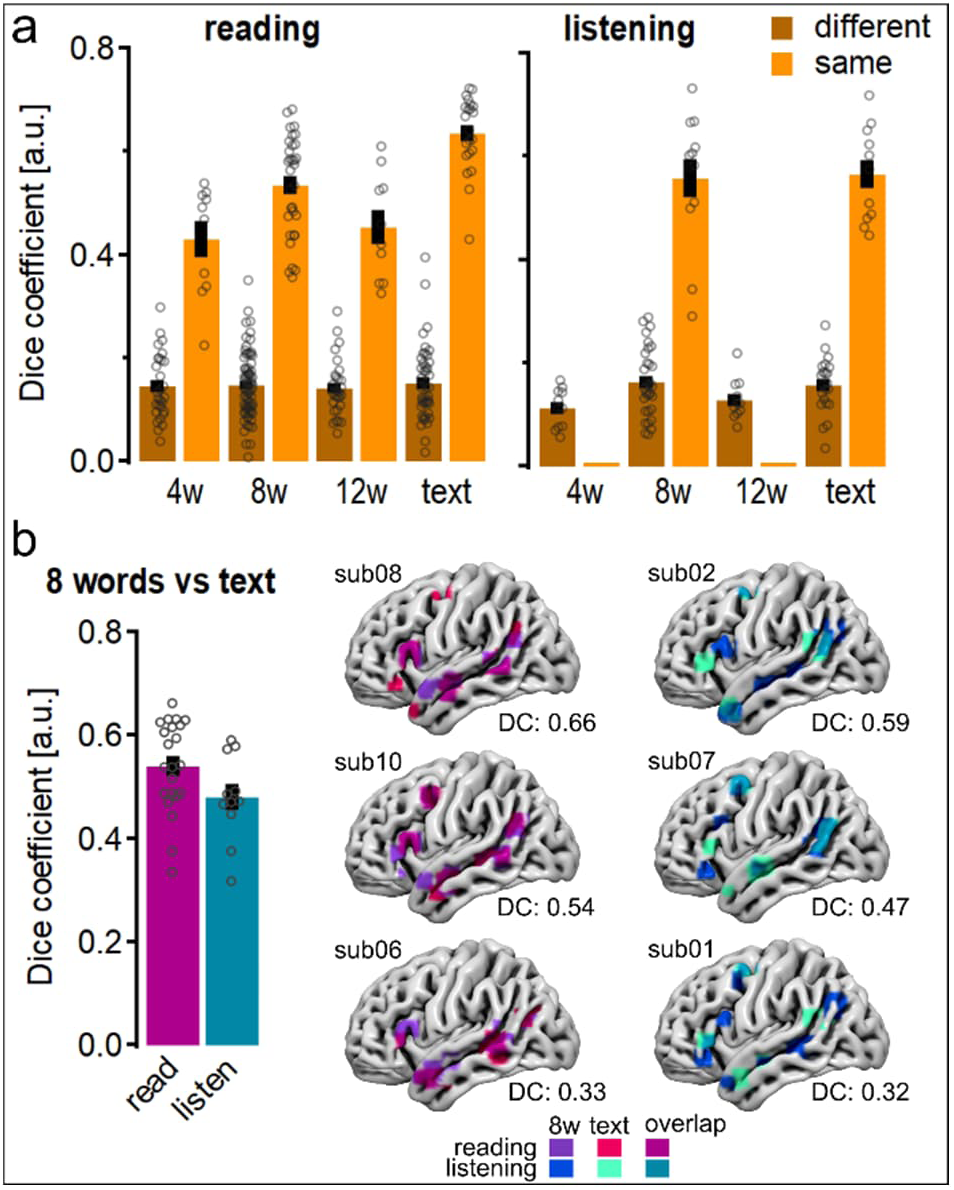
Spatial overlap of language fROIs defined by different conditions. (a) Dice coefficients were calculated for the left-hemisphere language fROIs within participants (orange) and between participants (brown bars) for the same task. (b) Within participants dice coefficients for the same task were compared between reading (purple) and listening (teal) language stimuli. Error bars represent the standard error of the mean. Brain-rendered surfaces show the overlap between 8-word sentences and text for reading and listening in representative participants with median dice coefficients, as well as in participants with the minimum and maximum values.

### Language network shows strongest responses to language stimuli

Across conditions, responses in the language network increased as stimuli became more syntactically connected, from short sentences to full texts, for both reading and listening, while activity in the MD and DM networks remained relatively stable (Fig. 5). To compare the relative engagement of the three networks we normalized effect sizes to the language network within each subject and condition, yielding the normalized values shown in Fig 6.

Modelling these relative effect sizes as a function of brain network (language, MD, DM), significantly improved model fit compared to the baseline model including modality, material, and their interaction (Model 1 vs. Model 2: *χ*²(2) = 81.6, *p* < 0.001), indicating robust differences in activation strength across brain networks. This effect did not vary by a network × material interaction (Model 2 vs. Model 3: *χ*²(6) = 6.5, *p* = 0.369), a network × modality interaction (Model 3 vs. Model 4: *χ*²(2) = 1.8 *p* = 0.409) or the three-way interaction (Model 4 vs. Model 5: *χ*²(6) = 3.6, *p* = 0.725), indicating no evidence for moderation by material or modality.

Bonferroni corrected pairwise comparisons of brain networks showed that relative effect sizes were larger in the language network than in the DM (*t*(425) = 2.7, *p* = 0.021) and MD networks (*t*(425) = 9.1, *p* < 0.001), and that the DM network also exceeded the MD network (*t*(425) = 6.4, *p* < 0.001) (Fig. 6). This pattern indicates that the language network is preferentially engaged by increasingly connected linguistic input, whereas the MD and DM networks do not track these changes as closely.

**Fig. 5.**
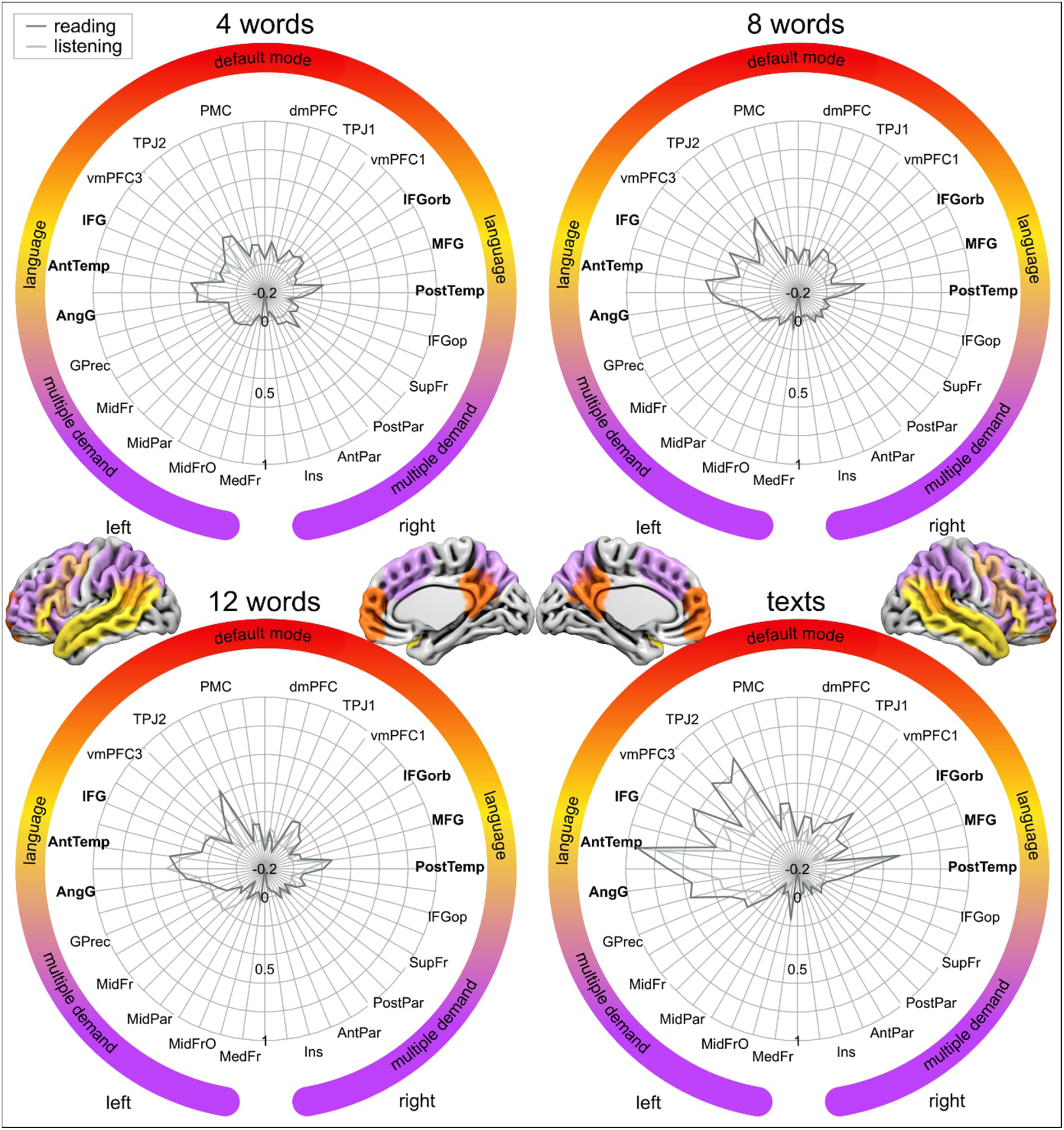
Language, default mode, and multiple demand network responses to increasing linguistic input length. Polar plots show average effect sizes for individual regions of interest (ROIs) in the language (yellow), default mode (red), and multiple demand (purple) networks during reading (gray line) and listening (light gray line) for four conditions: 4-word sentences, 8-word sentences, 12-word sentences, and short texts. ROI labels are split across the left and right sides of each polar plot to increase readability; labels appear on alternating ticks, with one set shown on the left side and the interleaved set shown on the right side, while the tick marks themselves are present on both sides. Cortical surface renderings illustrate the anatomical distribution of the three networks in the left and right hemispheres.

**Fig. 6.**
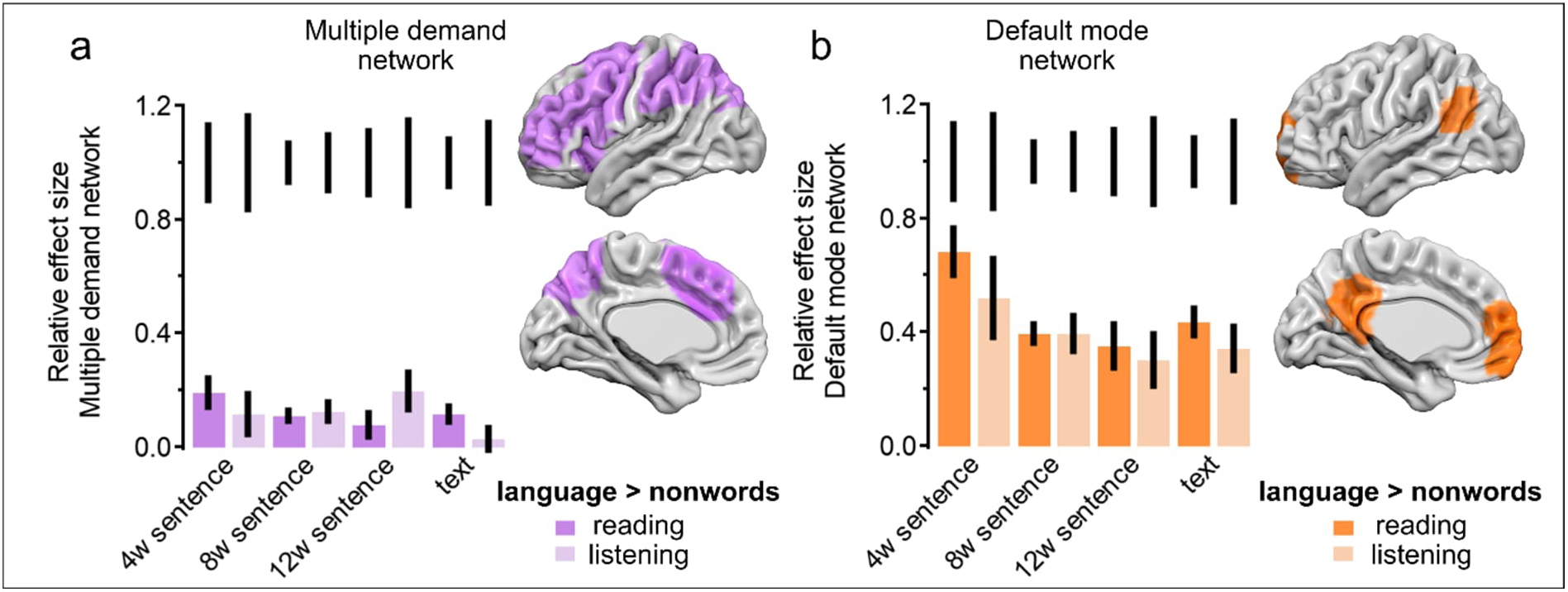
Neural responses in the multiple demand and default mode networks remained stable across input lengths. Bar plots showing the relative effect sizes in the multiple-demand (a) and default mode network (b), expressed as ratios to the average effect size in the language network across conditions. Error bars represent 95% CI. BOLD signal changes in both networks are positive but smaller than those in the language network, with significantly lower values in MD regions.

## DISCUSSION

Texts and longer, more connected sentences elicited stronger responses in the language network than do shorter sentences, suggesting that continuous, syntactically connected material is particularly suitable for clinical language mapping. This provides quantitative support for previous work showing that sentence-level and sentence-completion paradigms produce more extensive and reliable activation in classical language areas than simpler tasks (e.g. word lists or naming), and are therefore recommended in clinical guidelines [6,15]. Clinically, our findings support the use of sentence- and text-based comprehension tasks, rather than minimal sentence or word-list paradigms, to obtain robust, patient-tailored presurgical language maps in patients with no or only minor cognitive or language difficulties, particularly in left temporal and inferior frontal regions critical for language.

The graded increase from 4-word sentences to full texts in the language network is consistent with accounts that view sentence and discourse comprehension as hierarchical and context-dependent, with larger constituent structures requiring more combinatorial processing [44]. In line with this interpretation, Pallier et al. showed that activation in left inferior frontal and posterior temporal regions increases parametrically with the size of syntactic constituents, even when content words are replaced by pseudowords, indicating that these areas encode abstract hierarchical structure rather than mere word strings [44]. Our observation that longer, more connected materials (12-word sentences and texts) elicit stronger responses than short sentences particularly in left temporal and inferior frontal regions therefore likely reflects the incremental construction of larger syntactic and semantic constituents during continuous sentence and text processing.

A complementary line of evidence highlights the importance of semantic relatedness and conceptual integration for left inferior frontal engagement during sentence processing. An fMRI study manipulating syntactic complexity, processing phase, and semantic relatedness of sentence constituents showed that different subregions of the left inferior frontal gyrus are differentially sensitive to structural and semantic factors, consistent with a division of labor between syntactic operations and the integration of related concepts [45]. From a more abstract perspective, computational work on ‘gist’ extraction and topic-based semantic representations suggests that comprehension benefits when incoming input can be related to stored conceptual knowledge and used to predict and disambiguate upcoming material [46]. Taken together with our graded increase from short sentences to texts, and the relatively weaker activation of left inferior frontal regions for short sentences, these converging lines of evidence support the view that left inferior frontal cortex contributes not only to the construction of hierarchical structure but also to the integration of semantically related information and the extraction of higher-level conceptual gist over extended utterances.

The pronounced differences between within-subject and between-subject dice coefficients indicate high individual specificity of language maps, consistent with findings from precision fMRI [47,48], and current clinical recommendations for language mapping [6]. Although a common set of “core” regions emerges across participants, the fine-grained spatial patterns vary substantially, underscoring the need for patient-specific presurgical mapping rather than reliance on group templates. In our data, relatively high within-subject dice coefficients between sentence and text conditions further show that both materials engage largely overlapping language-sensitive regions, while the stronger responses to texts suggest that continuous, connected language provides a more robust signal for delineating each patient’s individual language network.

Although the modality-independent engagement of the left-hemispheric language network accords with common clinical practice and prior work showing broadly similar language maps for reading and listening [10,19,20], our data provide quantitative support for using either modality to localize language areas, which is particularly relevant when patients cannot perform one modality (e.g., due to visual or reading difficulties). At the same time, the stronger and more widespread effects for reading than listening, especially for texts, suggest that, when possible, visually presented connected language may yield more robust activation maps, with auditory texts serving as a viable alternative in patients for whom reading is not possible or reliable. However, the auditory modality is inherently more difficult to implement reliably as the scanner’s acoustic noise can mask or distort the stimulus, hearing sensitivity varies across individuals, and the necessary calibration of stimulus loudness to each participant introduces an additional source of variability that is largely avoided with visual presentation.

The stronger engagement of the language network relative to the MD and DM networks, together with the weak sensitivity of MD and DM to increases in linguistic connectedness, indicates that our paradigm predominantly probes language-selective cortex rather than domain-general control networks. This pattern is consistent with basic research showing a robust dissociation between language and MD systems and arguing that MD activations during language tasks often reflect task difficulty or executive demands rather than core linguistic processing [13,16,49]. Accordingly, our findings support the use of such paradigms for the specific identification of language-eloquent cortex in clinical settings, helping to reduce the risk of misattributing MD activity to essential language areas.

This study was conducted in healthy participants, limiting generalizability to patients [50]. Although each paradigm was successfully tested qualitatively in one patient, quantitative validation in a larger clinical sample remains necessary. Furthermore, clinical application of the subject-specific fROI approach requires validation against intraoperative mapping and postoperative outcomes. Finally, contributions from domain-general processes cannot be entirely excluded [17].

## CONCLUSION

Overall, our findings indicate that continuous sentence- and text-level paradigms based on ecologically valid, real-world language materials can provide a robust, language-selective, and patient-tailored approach to presurgical fMRI mapping of the left-hemispheric language network. Such connected, naturalistic language elicits strong activation in this network that is reliable across individuals yet sensitive to patient-specific anatomy, while predominantly recruiting language-selective rather than domain-general control regions, making these paradigms particularly suitable for presurgical language fMRI in patients who retain largely preserved cognitive and language abilities.

## Acknowledgements

We thank the study participants for their cooperation and patience.

## DECLARATIONS

### Ethical Standards

The study was approved by the Ethics committee of the University of Göttingen and was in accordance with the guidelines of the Declaration of Helsinki.

### Consent to participate

All participants gave written informed consent and obtained an expense allowance for taking part in this study.

## Abbreviations

AngG: angular gyrus
AntTemp: anterior temporal lobe
DCs: dice coefficients
DM: default mode network
fROI: subject-specific functional region of interest
IFG: inferior frontal gyrus
IFGorb: inferior frontal gyrus pars orbitalis
MD: multiple demand network
MFG: middle frontal gyrus
PostTemp: posterior temporal lobe

## REFERENCES

1. Sarubbo S, Duffau H. Connectomic evidences driving a functional approach in neuro-oncological surgery. J Neurosurg Sci. 2022;65:545–7. 10.23736/s0390-5616.21.05517-x

2. Ojemann G, Ojemann J, Lettich E, Berger M. Cortical language localization in left, dominant hemisphere: An electrical stimulation mapping investigation in 117 patients. J Neurosurg. 1989;71:316–26. 10.3171/jns.1989.71.3.0316

3. Urbach H, Mast H, Egger K, Mader I. Presurgical MR Imaging in Epilepsy. Clin Neuroradiol. 2015;25:151–5. 10.1007/s00062-015-0387-x

4. Duncan JS, Winston GP, Koepp MJ, Ourselin S. Brain imaging in the assessment for epilepsy surgery. Lancet Neurol. 2016;15:420–33. 10.1016/s1474-4422(15)00383-x

5. Sanvito F, Caverzasi E, Riva M, Jordan KM, Blasi V, Scifo P, et al. fMRI-Targeted High-Angular Resolution Diffusion MR Tractography to Identify Functional Language Tracts in Healthy Controls and Glioma Patients. Front Neurosci. 2020;14:225. 10.3389/fnins.2020.00225

6. Voets NL, Ashtari M, Beckmann CF, Benjamin CF, Benzinger T, Binder JR, et al. Consensus recommendations for clinical functional MRI applied to language mapping. Aperture Neuro. 2025;5. 10.52294/001c.128149

7. Zwart B de, Ruis C. An update on tests used for intraoperative monitoring of cognition during awake craniotomy. Acta Neurochir. 2024;166:204. 10.1007/s00701-024-06062-6

8. Ruis C. Monitoring cognition during awake brain surgery in adults: A systematic review. J Clin Exp Neuropsychol. 2018;40:1081–104. 10.1080/13803395.2018.1469602

9. Muir MT, Noll K, Patel R, Prinsloo S, Michener H, Kumar VA, et al. Toward data-driven surgical planning: multimodal mapping correlates of aphasic surgical deficits. J Neurosurg. 2026;144:324–35. 10.3171/2025.6.jns242301

10. Fedorenko E, Ivanova AA, Regev TI. The language network as a natural kind within the broader landscape of the human brain. Nat Rev Neurosci. 2024;1–24. 10.1038/s41583-024-00802-4

11. Binder JR. Functional Neuroradiology, Principles and Clinical Applications. In: Faro SH, Mohamed FB, editors. Functional Neuroradiology. Springer International Publishing AG; 2023. p. 565–92. 10.1007/978-3-031-10909-6_26

12. Black DF, Vachha B, Mian A, Faro SH, Maheshwari M, Sair HI, et al. American Society of Functional Neuroradiology–Recommended fMRI Paradigm Algorithms for Presurgical Language Assessment. Am J Neuroradiol. 2017;38:E65–73. 10.3174/ajnr.a5345

13. Mineroff Z, Blank IA, Mahowald K, Fedorenko E. A robust dissociation among the language, multiple demand, and default mode networks: Evidence from inter-region correlations in effect size. Neuropsychologia. 2018;119:501–11. 10.1016/j.neuropsychologia.2018.09.011

14. Fedorenko E, Hsieh P-J, Nieto-Castañón A, Whitfield-Gabrieli S, Kanwisher N. New Method for fMRI Investigations of Language: Defining ROIs Functionally in Individual Subjects. J Neurophysiol. 2010;104:1177–94. 10.1152/jn.00032.2010

15. Binder JR, Swanson SJ, Hammeke TA, Sabsevitz DS. A comparison of five fMRI protocols for mapping speech comprehension systems. Epilepsia. 2008;49:1980–97. 10.1111/j.1528-1167.2008.01683.x

16. Gao R, Cheung C, Siegelman M, Pongos ALA, Kean HH, Tanner A, et al. The language network responds robustly to sentences across tasks. Imaging Neurosci. 2026;4:IMAG.a.1283. 10.1162/imag.a.1283

17. Geranmayeh F, Wise RJS, Mehta A, Leech R. Overlapping Networks Engaged during Spoken Language Production and Its Cognitive Control. J Neurosci. 2014;34:8728–40. 10.1523/jneurosci.0428-14.2014

18. Fedorenko E, Blank IA. Broca’s Area Is Not a Natural Kind. Trends Cogn Sci. 2020;24:270–84. 10.1016/j.tics.2020.01.001

19. Braze D, Mencl WE, Tabor W, Pugh KR, Constable RT, Fulbright RK, et al. Unification of Sentence Processing via Ear and Eye: An fMRI Study. Cortex a J devoted study Nerv Syst Behav. 2011;47:416–31. 10.1016/j.cortex.2009.11.005

20. Cohen L, Salondy P, Pallier C, Dehaene S. How does inattention affect written and spoken language processing? Cortex. 2021;138:212–27. 10.1016/j.cortex.2021.02.007

21. Buchweitz A, Mason RA, Tomitch LMB, Just MA. Brain activation for reading and listening comprehension: An fMRI study of modality effects and individual differences in language comprehension. Psychol Neurosci. 2009;2:111–23. 10.3922/j.psns.2009.2.003

22. Oldfield RC. The assessment and analysis of handedness: The Edinburgh inventory. Neuropsychologia. 1971;9:97–113. 10.1016/0028-3932(71)90067-4

23. Keuleers E, Brysbaert M. Wuggy: A multilingual pseudoword generator. Behav Res Methods. 2010;42:627–33. 10.3758/brm.42.3.627

24. Jenkinson M, Beckmann CF, Behrens TEJ, Woolrich MW, Smith SM. FSL. NeuroImage. 2012;62:782–90. 10.1016/j.neuroimage.2011.09.015

25. Smith SM, Jenkinson M, Woolrich MW, Beckmann CF, Behrens TEJ, Johansen-Berg H, et al. Advances in functional and structural MR image analysis and implementation as FSL. NeuroImage. 2004;23:S208–19. 10.1016/j.neuroimage.2004.07.051

26. Woolrich MW, Jbabdi S, Patenaude B, Chappell M, Makni S, Behrens T, et al. Bayesian analysis of neuroimaging data in FSL. NeuroImage. 2009;45:S173–86. 10.1016/j.neuroimage.2008.10.055

27. Henschel L, Conjeti S, Estrada S, Diers K, Fischl B, Reuter M. FastSurfer - A fast and accurate deep learning based neuroimaging pipeline. NeuroImage. 2020;219:117012. 10.1016/j.neuroimage.2020.117012

28. Hagoort P, Indefrey P, Brown C, Herzog H, Steinmetz H, Seitz RJ. The Neural Circuitry Involved in the Reading of German Words and Pseudowords: A PET Study. J Cogn Neurosci. 1999;11:383–98. 10.1162/089892999563490

29. Friederici AD. The Brain Basis of Language Processing: From Structure to Function. Physiol Rev. 2011;91:1357–92. 10.1152/physrev.00006.2011

30. Hickok G, Poeppel D. The cortical organization of speech processing. Nat Rev Neurosci. 2007;8:393–402. 10.1038/nrn2113

31. Lipkin B, Tuckute G, Affourtit J, Small H, Mineroff Z, Kean H, et al. Probabilistic atlas for the language network based on precision fMRI data from >800 individuals. Sci Data. 2022;9:529. 10.1038/s41597-022-01645-3

32. Turker S, Kuhnke P, Eickhoff SB, Caspers S, Hartwigsen G. Cortical, Subcortical, and Cerebellar Contributions to Language Processing: A Meta-Analytic Review of 403 Neuroimaging Experiments. Psychol Bull. 2023;149:699–723. 10.1037/bul0000403

33. Fedorenko E, Duncan J, Kanwisher N. Broad domain generality in focal regions of frontal and parietal cortex. Proc National Acad Sci. 2013;110:16616–21. 10.1073/pnas.1315235110

34. Duncan J. The multiple-demand (MD) system of the primate brain: mental programs for intelligent behaviour. Trends Cogn Sci. 2010;14:172–9. 10.1016/j.tics.2010.01.004

35. Yeo BTT, Krienen FM, Sepulcre J, Sabuncu MR, Lashkari D, Hollinshead M, et al. The organization of the human cerebral cortex estimated by intrinsic functional connectivity. J Neurophysiol. 2011;106:1125–65. 10.1152/jn.00338.2011

36. Fox MD, Corbetta M, Snyder AZ, Vincent JL, Raichle ME. Spontaneous neuronal activity distinguishes human dorsal and ventral attention systems. Proc National Acad Sci. 2006;103:10046–51. 10.1073/pnas.0604187103

37. Buckner RL, Andrews-Hanna JR, Schacter DL. The brain’s default network: anatomy, function, and relevance to disease. Ann N York Acad Sci. 2008;1124:1–38. 10.1196/annals.1440.011

38. Kernbach JM, Yeo BTT, Smallwood J, Margulies DS, Schotten MT de, Walter H, et al. Subspecialization within default mode nodes characterized in 10,000 UK Biobank participants. Proc Natl Acad Sci. 2018;115:12295–300. 10.1073/pnas.1804876115

39. Buckner RL, DiNicola LM. The brain’s default network: updated anatomy, physiology and evolving insights. Nat Rev Neurosci. 2019;20:593–608. 10.1038/s41583-019-0212-7

40. Braga RM, Buckner RL. Parallel Interdigitated Distributed Networks within the Individual Estimated by Intrinsic Functional Connectivity. Neuron. 2017;95:457–471.e5. 10.1016/j.neuron.2017.06.038

41. Nieto-Castañón A, Fedorenko E. Subject-specific functional localizers increase sensitivity and functional resolution of multi-subject analyses. Neuroimage. 2012;63:1646–69. 10.1016/j.neuroimage.2012.06.065

42. Kriegeskorte N, Simmons WK, Bellgowan PSF, Baker CI. Circular analysis in systems neuroscience: the dangers of double dipping. Nat Neurosci. 2009;12:535–40. 10.1038/nn.2303

43. Dice LR. Measures of the Amount of Ecologic Association Between Species. Ecology. 1945;26:297–302. 10.2307/1932409

44. Pallier C, Devauchelle A-D, Dehaene S. Cortical representation of the constituent structure of sentences. Proc Natl Acad Sci. 2011;108:2522–7. 10.1073/pnas.1018711108

45. Newman SD, Ikuta T, Burns T. The effect of semantic relatedness on syntactic analysis: an fMRI study. Brain Lang. 2010;113:51–8. 10.1016/j.bandl.2010.02.001

46. Griffiths TL, Steyvers M, Tenenbaum JB. Topics in semantic representation. Psychol Rev. 2007;114:211–44. 10.1037/0033-295x.114.2.211

47. Wilson SM, Yen M, Eriksson DK. A n adaptive semantic matching paradigm for reliable and valid language mapping in individuals with aphasia. Hum Brain Mapp. 2018;39:3285–307. 10.1002/hbm.24077

48. Wilson SM, Bautista A, Yen M, Lauderdale S, Eriksson DK. Validity and reliability of four language mapping paradigms. NeuroImage: Clin. 2017;16:399–408. 10.1016/j.nicl.2016.03.015

49. Diachek E, Blank I, Siegelman M, Affourtit J, Fedorenko E. The Domain-General Multiple Demand (MD) Network Does Not Support Core Aspects of Language Comprehension: A Large-Scale fMRI Investigation. J Neurosci. 2020;40:4536–50. 10.1523/jneurosci.2036-19.2020

50. Pak RW, Hadjiabadi DH, Senarathna J, Agarwal S, Thakor NV, Pillai JJ, et al. Implications of neurovascular uncoupling in functional magnetic resonance imaging (fMRI) of brain tumors. J Cereb Blood Flow Metab. 2017;37:3475–87. 10.1177/0271678x17707398

