## Supplementary for "The language network responds stronger to connected than to unconnected sentences"

### Supplementary material

**The language network responds stronger to connected than to unconnected sentences**

**Supplementary Table 1 Subjects and Conditions**

| *N* | Modality | | Material | | | | |
|---|---|---|---|---|---|---|---|
| | Auditory | Visual | 4w, 2 runs | 8w, 2 runs | 12w, 2 runs | Texts, 2 runs | 4w, 8w, 12w, 1 run each |
| 11 | x | x | | x | | x | |
| 10 | | x | x | x | x | | |
| 10 | x | x | | | | | x |
| 11 | | x | | x | | x | |
| 1 | x | | x | x | x | | |

**Supplementary Table 2 Results of the pairwise comparison of effect sizes averaged across reading and listening**

| Pair | Estimate | SE | df | t-ratio | p-value |
|---|---|---|---|---|---|
| Left IFGorb | | | | | |
| Texts - 12-word | 0.291 | 0.048 | 882 | 6.067 | **< 0.001** |
| Texts - 8-word | 0.232 | 0.039 | 882 | 5.877 | **< 0.001** |
| Texts - 4-word | 0.425 | 0.048 | 882 | 8.864 | **< 0.001** |
| 12-words - 8-word | 0.059 | 0.040 | 882 | 1.465 | 0.860 |
| 12-words - 4-word | 0.134 | 0.046 | 882 | 2.928 | **0.021** |
| 8-words - 4-word | 0.193 | 0.040 | 882 | 4.789 | **< 0.001** |
| Left IFG | | | | | |
| Texts - 12-word | 0.208 | 0.048 | 882 | 4.338 | **< 0.001** |
| Texts - 8-word | 0.207 | 0.039 | 882 | 5.243 | **< 0.001** |
| Texts - 4-word | 0.315 | 0.048 | 882 | 6.573 | **< 0.001** |

| | | | | | |
|---|---|---|---|---|---|
| 12-words - 8-word | -0.001 | 0.040 | 882 | -0.030 | 1.0 |
| 12-words - 4-word | 0.107 | 0.046 | 882 | 2.340 | 0.117 |
| 8-words - 4-word | 0.108 | 0.040 | 882 | 2.687 | **0.044** |
| Left MFG | | | | | |
| Texts - 12-word | 0.083 | 0.048 | 882 | 1.735 | 0.499 |
| Texts - 8-word | 0.121 | 0.039 | 882 | 3.078 | **0.013** |
| Texts - 4-word | 0.224 | 0.048 | 882 | 4.665 | < **0.001** |
| 12-words - 8-word | 0.038 | 0.040 | 882 | 0.947 | 1.0 |
| 12-words - 4-word | 0.141 | 0.046 | 882 | 3.067 | **0.013** |
| 8-words - 4-word | 0.102 | 0.040 | 882 | 2.535 | 0.069 |
| Left Ang | | | | | |
| Texts - 12-word | 0.145 | 0.048 | 882 | 3.029 | **0.015** |
| Texts - 8-word | 0.169 | 0.039 | 882 | 4.287 | < **0.001** |
| Texts - 4-word | 0.223 | 0.048 | 882 | 4.640 | < **0.001** |
| 12-words - 8-word | 0.024 | 0.040 | 882 | 0.591 | 1.0 |
| 12-words - 4-word | 0.077 | 0.046 | 882 | 1.686 | 0.553 |
| 8-words - 4-word | 0.053 | 0.040 | 882 | 1.323 | 1.0 |
| Left AntTemp | | | | | |
| Texts - 12-word | 0.431 | 0.048 | 882 | 8.981 | < **0.001** |
| Texts - 8-word | 0.455 | 0.039 | 882 | 11.532 | < **0.001** |
| Texts - 4-word | 0.563 | 0.048 | 882 | 11.749 | < **0.001** |
| 12-words - 8-word | 0.024 | 0.040 | 882 | 0.600 | 1.0 |
| 12-words - 4-word | 0.133 | 0.046 | 882 | 2.898 | **0.023** |
| 8-words - 4-word | 0.109 | 0.040 | 882 | 2.690 | **0.044** |
| Left PostTemp | | | | | |
| Texts - 12-word | 0.197 | 0.048 | 882 | 4.113 | < **0.001** |
| Texts - 8-word | 0.240 | 0.039 | 882 | 6.082 | < **0.001** |
| Texts - 4-word | 0.357 | 0.048 | 882 | 7.454 | < **0.001** |
| 12-words - 8-word | 0.043 | 0.040 | 882 | 1.057 | 1.0 |
| 12-words - 4-word | 0.160 | 0.046 | 882 | 3.497 | **0.003** |
| 8-words - 4-word | 0.118 | 0.040 | 882 | 2.913 | **0.022** |

**Supplementary Table 3 Results of the pairwise comparison of effect sizes in visual modality**

| Pair | Estimate | SE | df | t-ratio | p-value |
|---|---|---|---|---|---|
| Left IFGorb | | | | | |
| Texts - 12-word | 0.379 | 0.058 | 864 | 6.562 | < **0.001** |
| Texts - 8-word | 0.276 | 0.048 | 864 | 5.761 | < **0.001** |
| Texts - 4-word | 0.480 | 0.058 | 864 | 8.298 | < **0.001** |

| | | | | | |
|---|---|---|---|---|---|
| 12-words - 8-word | -0.104 | 0.049 | 864 | -2.098 | 0.217 |
| 12-words - 4-word | 0.100 | 0.057 | 864 | 1.770 | 0.462 |
| 8-words - 4-word | 0.204 | 0.049 | 864 | 4.127 | **< 0.001** |
| Left IFG | | | | | |
| Texts - 12-word | 0.289 | 0.058 | 864 | 5.003 | **< 0.001** |
| Texts - 8-word | 0.298 | 0.048 | 864 | 6.236 | **< 0.001** |
| Texts - 4-word | 0.408 | 0.058 | 864 | 7.055 | **< 0.001** |
| 12-words - 8-word | 0.009 | 0.049 | 864 | 0.183 | 1.0 |
| 12-words - 4-word | 0.119 | 0.057 | 864 | 2.092 | 0.220 |
| 8-words - 4-word | 0.110 | 0.049 | 864 | 2.214 | 0.162 |
| Left MFG | | | | | |
| Texts - 12-word | 0.110 | 0.058 | 864 | 1.910 | 0.338 |
| Texts - 8-word | 0.153 | 0.048 | 864 | 3.190 | **0.009** |
| Texts - 4-word | 0.272 | 0.058 | 864 | 4.710 | **< 0.001** |
| 12-words - 8-word | 0.042 | 0.049 | 864 | 0.852 | 1.0 |
| 12-words - 4-word | 0.162 | 0.057 | 864 | 2.855 | **0.026** |
| 8-words - 4-word | 0.120 | 0.049 | 864 | 2.420 | 0.094 |
| Left Ang | | | | | |
| Texts - 12-word | 0.176 | 0.058 | 864 | 3.049 | **0.014** |
| Texts - 8-word | 0.202 | 0.048 | 864 | 4.217 | **< 0.001** |
| Texts - 4-word | 0.243 | 0.058 | 864 | 4.207 | **< 0.001** |
| 12-words - 8-word | 0.025 | 0.049 | 864 | 0.514 | 1.0 |
| 12-words - 4-word | 0.067 | 0.057 | 864 | 1.181 | 1.0 |
| 8-words - 4-word | 0.042 | 0.049 | 864 | 0.840 | 1.0 |
| Left AntTemp | | | | | |
| Texts - 12-word | 0.438 | 0.058 | 864 | 7.580 | **< 0.001** |
| Texts - 8-word | 0.467 | 0.048 | 864 | 9.774 | **< 0.001** |
| Texts - 4-word | 0.587 | 0.058 | 864 | 10.158 | **< 0.001** |
| 12-words - 8-word | 0.029 | 0.049 | 864 | 0.593 | 1.0 |
| 12-words - 4-word | 0.149 | 0.057 | 864 | 2.629 | 0.052 |
| 8-words - 4-word | 0.120 | 0.049 | 864 | 2.420 | 0.094 |
| Left PostTemp | | | | | |
| Texts - 12-word | 0.267 | 0.058 | 864 | 4.626 | **< 0.001** |
| Texts - 8-word | 0.282 | 0.048 | 864 | 5.901 | **< 0.001** |
| Texts - 4-word | 0.412 | 0.058 | 864 | 7.136 | **< 0.001** |
| 12-words - 8-word | 0.015 | 0.049 | 864 | 0.300 | 1.0 |
| 12-words - 4-word | 0.145 | 0.057 | 864 | 2.560 | 0.064 |
| 8-words - 4-word | 0.130 | 0.049 | 864 | 2.633 | 0.052 |

**Supplementary Table 4 Results of the pairwise comparison of effect sizes in auditory modality**

| Pair | Estimate | SE | df | t-ratio | p-value |
|---|---|---|---|---|---|
| Left IFGorb | | | | | |
| Texts - 12-word | 0.118 | 0.079 | 864 | 1.495 | 0.812 |
| Texts - 8-word | 0.142 | 0.067 | 864 | 2.126 | 0.203 |
| Texts - 4-word | 0.313 | 0.079 | 864 | 3.981 | **< 0.001** |
| 12-words - 8-word | 0.025 | 0.067 | 864 | 0.367 | 1.0 |
| 12-words - 4-word | 0.196 | 0.076 | 864 | 2.559 | 0.064 |
| 8-words - 4-word | 0.171 | 0.067 | 864 | 2.559 | 0.064 |
| Left IFG | | | | | |
| Texts - 12-word | 0.043 | 0.079 | 864 | 0.548 | 1.0 |
| Texts - 8-word | 0.024 | 0.067 | 864 | 0.353 | 1.0 |
| Texts - 4-word | 0.130 | 0.079 | 864 | 1.647 | 0.599 |
| 12-words - 8-word | -0.020 | 0.067 | 864 | -0.292 | 1.0 |
| 12-words - 4-word | 0.086 | 0.076 | 864 | 1.131 | 1.0 |
| 8-words - 4-word | 0.106 | 0.067 | 864 | 1.586 | 0.679 |
| Left MFG | | | | | |
| Texts - 12-word | 0.024 | 0.079 | 864 | 0.308 | 1.0 |
| Texts - 8-word | 0.057 | 0.067 | 864 | 0.857 | 1.0 |
| Texts - 4-word | 0.126 | 0.079 | 864 | 1.601 | 0.658 |
| 12-words - 8-word | 0.033 | 0.067 | 864 | 0.495 | 1.0 |
| 12-words - 4-word | 0.102 | 0.076 | 864 | 1.332 | 1.0 |
| 8-words - 4-word | 0.069 | 0.067 | 864 | 1.028 | 1.0 |
| Left Ang | | | | | |
| Texts - 12-word | 0.080 | 0.079 | 864 | 1.019 | 1.0 |
| Texts - 8-word | 0.103 | 0.067 | 864 | 1.535 | 0.750 |
| Texts - 4-word | 0.176 | 0.079 | 864 | 2.238 | 0.153 |
| 12-words - 8-word | 0.022 | 0.067 | 864 | 0.336 | 1.0 |
| 12-words - 4-word | 0.096 | 0.076 | 864 | 1.255 | 1.0 |
| 8-words - 4-word | 0.073 | 0.067 | 864 | 1.099 | 1.0 |
| Left AntTemp | | | | | |
| Texts - 12-word | 0.411 | 0.079 | 864 | 5.219 | **< 0.001** |
| Texts - 8-word | 0.428 | 0.067 | 864 | 6.399 | **< 0.001** |
| Texts - 4-word | 0.514 | 0.079 | 864 | 6.531 | **< 0.001** |
| 12-words - 8-word | 0.017 | 0.067 | 864 | 0.256 | 1.0 |
| 12-words - 4-word | 0.103 | 0.076 | 864 | 1.350 | 1.0 |
| 8-words - 4-word | 0.086 | 0.067 | 864 | 1.288 | 1.0 |
| Left PostTemp | | | | | |
| Texts - 12-word | 0.058 | 0.079 | 864 | 0.733 | 1.0 |
| Texts - 8-word | 0.153 | 0.067 | 864 | 2.291 | 0.133 |
| Texts - 4-word | 0.245 | 0.079 | 864 | 3.119 | **0.011** |

| | | | | | |
|---|---|---|---|---|---|
| 12-words - 8-word | 0.095 | 0.067 | 864 | 1.428 | 0.922 |
| 12-words - 4-word | 0.188 | 0.076 | 864 | 2.456 | 0.086 |
| 8-words - 4-word | 0.092 | 0.067 | 864 | 1.380 | 1.0 |

**Supplementary Table 5 Results of the pairwise comparison of DCs**

| **Material** | **Pair** | **Estimate** | **SE** | **df** | **t-ratio** | **p-value** |
|---|---|---|---|---|---|---|
| 4-word | Within – between subject | 0,301 | 0,026 | 201 | 11,61 | **<0.001** |
| 8-word | Within – between subject | 0,388 | 0,014 | 201 | 27,891 | **<0.001** |
| 12-word | Within – between subject | 0,319 | 0,026 | 201 | 12,276 | **<0.001** |
| Texts | Within – between subject | 0,459 | 0,017 | 201 | 26,911 | **<0.001** |
